# CooccurrenceAffinity: An R package for computing a novel metric of affinity in co-occurrence data that corrects for pervasive errors in traditional indices

**DOI:** 10.1101/2022.11.01.514801

**Authors:** Kumar P. Mainali, Eric Slud

## Abstract

1. Analysis of co-occurrence data in ecology and biogeography with traditional indices has led to many problems. In our recent study (Mainali, Slud, Singer, & Fagan, 2022), we revealed the source of the problem that makes the traditional indices fundamentally flawed and completely unreliable, and we further developed a novel metric of association, alpha, with complete formulation of the null distribution for estimating the mechanism of affinity. We also developed the maximum likelihood estimate (MLE) of alpha in our prior study.
2. Here, we introduce the `CooccurrenceAffinity` R package that computes alpha MLE. We provide functions to perform the analysis based on a 2×2 contingency table of occurrence/co-occurrence counts as well as a m×n presence-absence matrix (e.g., species by site matrix). The flexibility of the function allows a user to compute the alpha MLE for entity pairs on matrix columns based on presence-absence states recorded in the matrix rows, or for entities on matrix rows based on presence-absence recorded in columns. We also provide functions for plotting the computed indices.
3. As novel components of this software paper not reported in the original study, we present theoretical discussions about median interval and four types of confidence intervals; we further develop functions (a) to compute those intervals, (b) to evaluate their true coverage probability of enclosing the population parameter, and (c) to generate figures.
4. `CooccurrenceAffinity` is a practical and efficient R package with user-friendly functions for end-to-end analysis and plotting of co-occurrence data in various formats, making it possible to compute the recently developed metric of alpha MLE as well as its median and confidence intervals introduced in this paper. The package supplements its main output of the novel metric of association with the three most common traditional indices of association in co-occurrence data: Jaccard, Sørensen–Dice, and Simpson.

## INTRODUCTION

Cooccurrence data are analyzed to estimate similarities among entities in many disciplines including biogeography (de Forges, Koslow, & Poore, 2000; Blowes et al., 2019), biodiversity (Chase & Leibold, 2002; Dornelas et al., 2014), ecology (Gotelli & McCabe, 2002), epidemiology (Cornfield, 1956), evolution (Plata, Henry, & Vitkup, 2015), and neuroscience (Crossley et al., 2013). Over a century’s quantitative development has resulted in ~80 metrics of association in co-occurrence data (Keil, 2019), of which the most popular metrics are Jaccard, Sørensen–Dice, and Simpson (Hohn, 2018). In a recent study (Mainali et al., 2022), we showed that these indices of association suffer from a fundamental statistical flaw, making them completely unreliable as a measure of association. For example, we show situations when the same value of Jaccard’s Index (e.g., 0.6) indicates strong negative and strong positive association (see Fig. 1D in (Mainali et al., 2022)). This lack of reliability of the traditional indices has been reflected in myriads of publications about beta diversity; see (Mainali et al., 2022) for a detailed account of problems of traditional indices from the viewpoint of statistical theory as well as of ecological practice.

**Figure 1.**
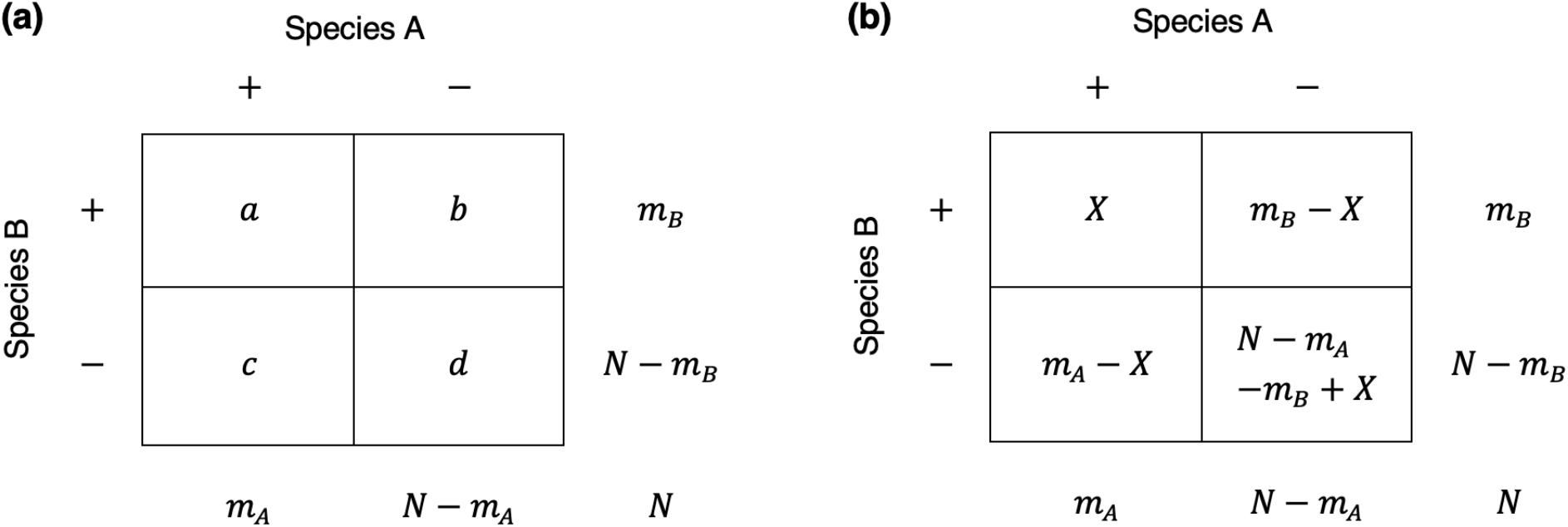
Two ways of displaying a 2×2 table of co-occurrences and occurrences of two entities with fixed margins. An example of distribution of Species A and B occupying *m_A_* and *m_B_* sites, respectively, out of a total *N* sites is presented here. The number of sites where both species cooccur (top-left quadrant), where only Species A exists (bottom-left quadrant), where only Species B exists (top-right quadrant), and where none of the species exist (bottom-right quadrant) are typically presented in popular scientific literature with a set of notations (Fig. 1a) but are often presented in statistical literature with a different set of notations (Fig. 1b).

We recently resolved the challenges of traditional indices by developing a reliable, meaningful and interpretable parameter of association in binary cooccurrence data. We developed this novel parameter, named alpha (*α*), from solid statistical theory and presented a complete mathematical formulation of its probability distribution in (Mainali et al., 2022). The purpose of this package is fivefold: (1) compute and display point estimate (Maximum Likelihood Estimate) of *α* or *log-affinity* as advanced in (Mainali et al., 2022), (2) present the mathematical basis of median interval and four types of confidence intervals, (3) compute the aforementioned five types of intervals, (4) develop a pipeline to analyze data that comes in standard 2×2 contingency table of co-occurrences and occurrences as well as entities by elements matrix (e.g., species by site matrix) of presence-absence states, and (5) generate plots with easy customization of graphical elements.

## MODELS, PARAMETERS AND INTERVALS

Imagine an ecological example involving distribution of two species on a grid of *N* sites in a specified locale. The occurrence data are typically given for each species pair in a 2×2 table with fixed margins (Fig. 1a,b) where *m_A_*, *m_B_* are the respective numbers of ecologic sites occupied by species A and B (prevalence of Species A = *m_A_/N*). The figure shows two types of notations used to represent the number of sites that contain both species, only species A, only species B and none of them.

The probability model for these data is a ‘balls-in-boxes’ or Urn Model. It encompasses both the null-hypothetical model of random assortment subject to given prevalences *m_A_*, *m_B_* and numbers *N* of sites, and also a single-parameter model expressing a quantitative affinity relating the probabilities with which the two species occupy each of the sites.

### The parameter

The model assumes that species A occupies *m_A_* sites out of the *N* equiprobably, that is, with each set of *m_A_* sites just as likely to be occupied as every other. Then species B occupies sites by choosing them independently of one another, with probability *p*_1_ if the site is already occupied by species A and with probability *p*_2_ if the site is not occupied (Fig. 2a). The random variable *X* of interest is the number of sites occupied by both species A and B. The probability distribution of *X* is computed conditionally given that the total number of sites occupied by species B is fixed at *m_B_*. This probability distribution is called Extended Hypergeometric (Harkness, 1965) or Noncentral Hypergeometric (Fisher, 1935). It turns out that this distribution would be the same if the roles of A, B were reversed (and the *m_B_* sites occupied by B were picked first), and the distribution depends on the probabilities *p*_1_, *p*_2_ only through the single log odds-ratio parameter

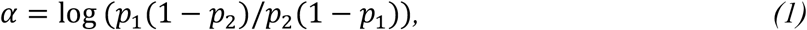

and has probability mass function (conditional on *m_A_*, *m_B_*, *N*) for max(*m_A_* + *m_B_*-N, 0) ≤ k ≤ min(*m_A_*,, *m_B_*) given by

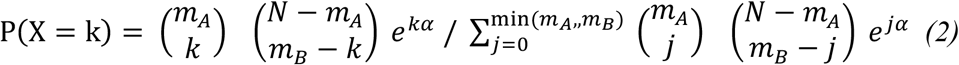

**Figure 2.**
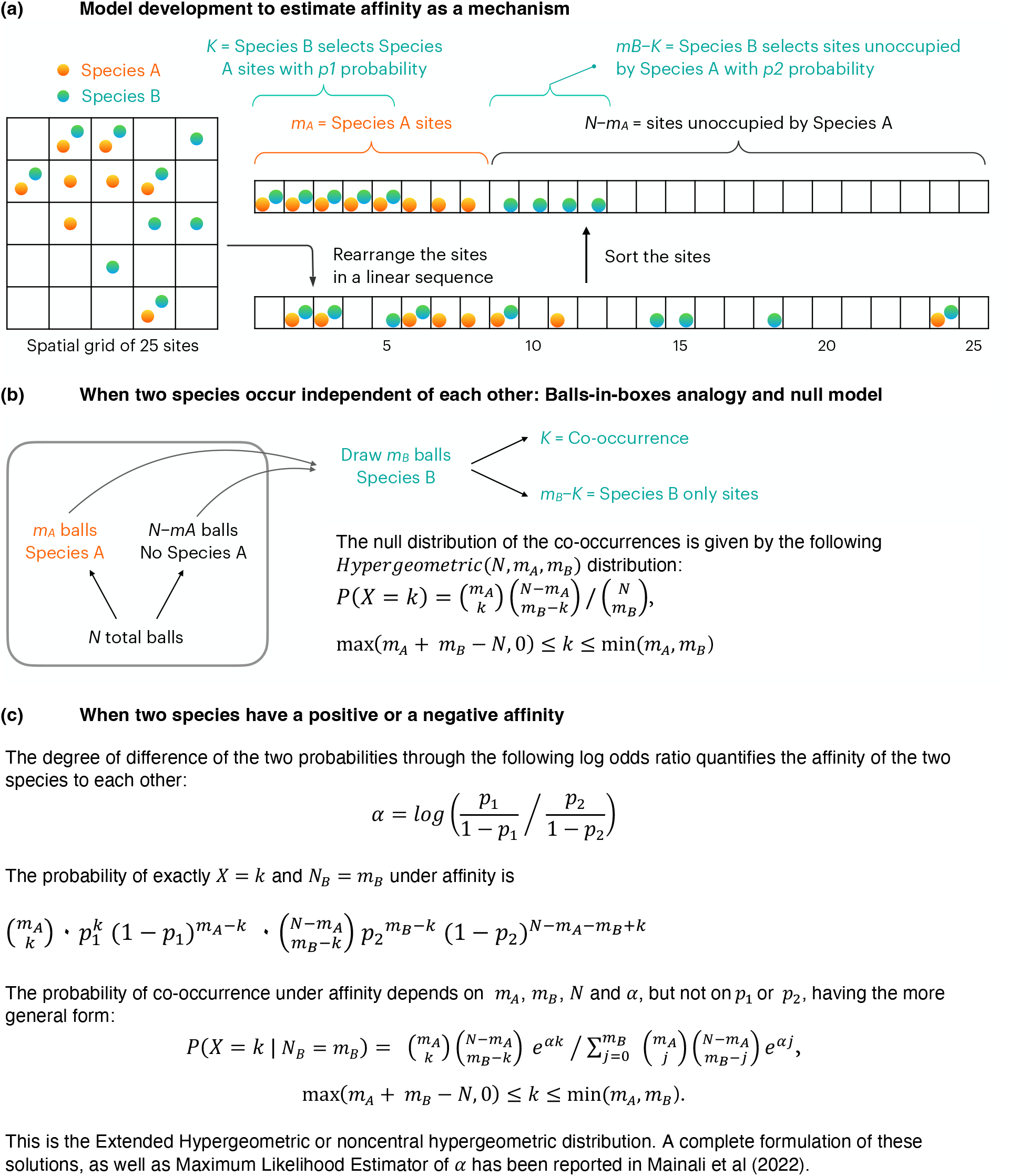
Model development for estimating affinity between two species as a mechanism. The three stages of the parameter development show the structure of affinity that characterizes the mechanism of affinity with mathematical logic (a). We next present the structure of the null model for the situation of zero affinity as Hypergeometric distribution (b) and extend this null for the case of nonzero affinity showing that it takes the form of Extended Hypergeometric distribution, leading to an estimate of alpha out of 2×2 table.

The parameter α defined in equation (1) is called log odds-ratio in standard statistical presentations of categorical data, and is ¼ times the row-column interaction parameter in standard parameterizations of *loglinear models* (Bishop, Fienberg, & Holland, 1975) on a 2×2 table. This unknown statistical parameter that quantifies the degree to which species A, B appear together – or that quantifies the affinity between any two entities based on their cooccurrence pattern – was recently advanced in the Ecology context in (Mainali et al., 2022). The statistical estimate of this parameter by maximum likelihood is called the alpha hat in (Mainali et al., 2022) and is shown there to be much preferable to standard ecologic indices of species-pair association or beta diversity because it quantifies dependence in a way insensitive to and interpretable apart from prevalences.

The complete mathematical formulation of alpha metric is available in (Mainali et al., 2022). In Fig. 2, we present a semi-graphical summary of alpha metric development. For the scenario of two species described above, the log odds ratio (the log of ratio of relative preference of Species B to species A sites to the relative preference of Species B to non-Species A sites) is the most intuitive and mechanistic expression of the true affinity between the two species (Fig. 2a). When the two species have zero affinity to each other, the arrangement in Fig. 2a naturally agrees with the balls-in-boxes description of the Hypergeometric null distribution (Fig. 2b). The mathematical form of the null distribution was recognized within ecological literature recently by (Veech, 2013) and (Griffith, Veech, Marsh, & others, 2016). When the two species have some affinity (positive or negative) to each other, their selective preference modifies the Hypergeometric equation as shown in Fig. 2c, to yield the Extended Hypergeometric (Fig. 2c). A beauty of this solution is that the log odds ratio originally computed with preferences of the two species (eq. 1) emerges out of Extended Hypergeometric distribution by being completely free from these preferences (*p*_1_, *p*_2_).

### The Maximum Likelihood Estimator

The Maximum Likelihood Estimator (MLE) 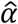 is computed from data (*X, m_A_*, *m_B_*, *N*), by numerically maximizing the right-hand side of equation (2) with *k* replaced by the observed *X* value. It has all the usual favorable properties of MLE’s (including approximate normal distribution when the sample sizes are large, which means here that min(*m_A_*, *m_B_*, *N* – *m_A_*, *N* – *m_B_*) is large), but is also meaningful when samples are not large. Along with the MLE, the package computes confidence intervals for the unknown parameter that have coverage probabilities reliably close to their nominal coverage probabilities apart from very extreme small-sample cases, as we will see below.

When *k* = max(*m_A_* + *m_B_* – *N*, 0) or *k* = min(*m_A_*, *m_B_*), the extended-real-valued maximizer of (2) occurs respectively at *α* = –∞ or +∞. To avoid these infinite values, we follow a Bayesian argument sketched in (Mainali et al., 2022) and modify the definition of the MLE 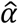 *only in the case where X is one of these extreme endpoints* respectively to ∓log(2*N*^2^).

### Advances: The median interval

When sample sizes are small the discreteness of the probability distribution (2) is an obstacle to precise estimation of α or confidence intervals covering it. One aspect of this discreteness is that for each q ϵ (0, 1) and each *k* satisfying max(*m_A_* + *m_B_* – *N*, 0) < *k* < min(*m_A_*, *m_B_*), there is not a single value but rather an interval of values for *α* that are compatible with *k* being a *q*’th quantile of (2), defined by the simultaneous inequalities

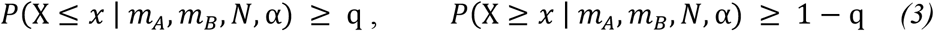

Denote by *F*(*x, α*) = *P*(*X* ≤ *x* | *m_A_, m_B_, N*, α) = *F*(*x, m_A_, m_B_, N*, α) the Extended Hypergeometric distribution function for the random variable *X* which is the upper-left entry of a 2×2 table with fixed first row-sum *m_A_*, first column-sum *m_B_*, table-total *N*, and odds-ratio parameter *e^α^*. It is easy to verify from (2) that *F*(x, α) is strictly monotonically decreasing in α, converging to 0 when *α* → ∞ and to 1 when α → −∞ for max(*m_A_* + *m_B_* – *N*, 0) < *x* < min(*m_A_*, *m_B_*). For such *x*, we denote by *F*(*x*, ·)^−1^ the inverse function with respect to *α*, from which it follows immediately that the largest open interval of *α* values satisfying (4) is (*F*(*x* – 1, ·)^−1^(*p*), *F*(*x*, ·)^−1^(*p*)), where *F*(*z*, ·)^−1^(*p*) is defined equal to that value *a* for which *F*(*z, a*) = *p*. All quantities *F*(*x, a*) computed in the `CooccurrenceAffinity` package are obtained from the function `pFNCHypergeo` in the R package `BiasedUrn`. Note that, for reasons of numerical accuracy and convergence of the functions in the `BiasedUrn` package, all absolute values of 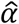 are capped at the largest value less than or equal to 10 for which the `BiasedUrn` functions work, usually 10 or close to it.

The interval described in the previous paragraph for *x* = *X* and *p* = 1/2 will be called the *median interval* for *α* based on *X*. This interval in our experience always contains the MLE of 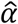 (an empirical finding, always found to hold numerically but not yet proved mathematically), and its length quantifies the discreteness in the Extended Hypergeometric distribution and thus the ambiguity of 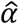.

### Advances: Four types of confidence interval

Next, we discuss the computation of the (two-sided equal-tailed) Confidence Interval for *α* based on the data (*X, m_A_, m_B_, N*). The concept of “test-based confidence interval” is central to explaining the confidence intervals for log-affinity parameter computed in this package. The idea is explained well in [(Casella & Berger, 2002), Sec.9.2.1] under the heading “Inverting a Test” and is familiar – sometimes by other names – from the history of confidence intervals for an unknown binomial proportion. Essentially the same idea is standard in defining so-called ‘exact’ confidence intervals for an unknown scalar parameter that is monotonically related to the distribution function for a discrete random variable. This construction applies to the parameter for Binomial, Poisson, Geometric, and Negative Binomial random variables, as well as for the log odds-ratio or affinity parameter *α* of the Extended Hypergeometric (Harkness, 1965) random variable, but we describe it in detail only in the affinity case.

The common feature shared by all of these discrete (integer-valued) random variables *X* with parameter *θ* is the Monotone Likelihood Ratio (MLR) property [(Casella & Berger, 2002), p.391] that states that the ratio *p*(*x*, *θ*_1_)/*p*(*x*, *θ*_2_) for fixed *θ*_1_ < *θ*_2_ is a monotonic function of integers *x*. This property, which is also a consequence of the natural-exponential-family property shared by these random variables, implies that the rejection regions for optimal (Neyman-Pearson) hypothesis tests of *H*_0_:*θ* ≤ *θ*_0_ versus *H*_1_:*θ* > *θ*_0_ or of *H*′_0_: *θ* ≥ *θ*_0_ versus *H′*_2_: *θ* < *θ*_0_ are respectively intervals {*x*: *x* ≥ *k*_2_} or {*x*: *x* ≤ *k*}. This is (part of) the Karlin-Rubin Theorem [(Casella & Berger, 2002), Theorem 8.3.17). In view of this discussion along with the monotone decrease of *F*(*x, α*) with respect to *α*, a natural equal-tailed test with significance level *γ* for the point null-hypothesis *H*_0_:*α* = *α*_0_ versus the two-sided alternative *H_A_*: *α* ≠ *α*_0_, would reject when *X* is too large or small, i.e., when *X* ≤ *k*_1_ or *X* ≥ *k*_2_ where the cut-offs *k*_1_, *k*_2_ are respectively determined as

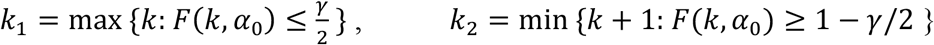

Thus, the acceptance region (complement of rejection region) for the test is the set of x’s for which

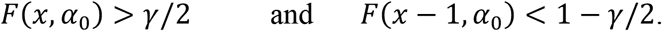

#### Clopper-Pearson type Confidence Interval

In terms of this family of hypothesis tests, the idea of the “test-based” or “inverted” confidence intervals for *α* based on *X* is to take as confidence interval *CI*(*X*, 1 – *γ*) of level 1 – *γ* the set of parameter values *α*_0_ for which the corresponding hypothesis tests would accept *H*_0_:*α* = *α*_0_. The idea of the test-based interval is that the Confidence Interval is the set of *α*_0_ values compatible with the data *X*.

This confidence set is an interval because *F*(*x*, *α*) is strictly decreasing with respect to *α*. Suppose that *x* is a possible value of the Extended Hypergeometric random variable, which is equivalent to saying that max(*m_A_* + *m_B_* – *N*, 0) ≤ *x* ≤ min(*m_A_*, *m_B_*). Recall that *F*(*x*, ·)^−1^(*r*) for 0 < *r* < 1 is the unique value of *α* ϵ (−∞, ∞) for which *F*(*x*, *α*) = *F*(*x, m_A_, m_B_, N*, α) = *r*. Then the acceptance region for the test *H*_0_:*α* = *α*_0_ above can equivalently be written

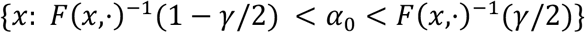

or as the ***Clopper-Pearson type Confidence Interval (CI.CP)***

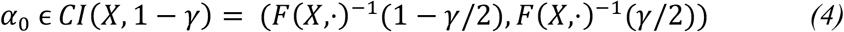

The statement that the test *H*_0_:*α* = *α*_0_ has significance level *γ* is exactly the same as the statement that the probability of *α*_0_ ϵ *CI*(*X*, 1 – *γ*) is at least 1 – *γ*.

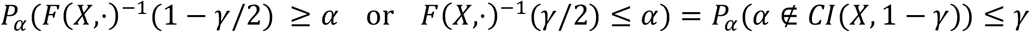

The BiasedUrn function pFNCHypergeo computes the function *F*(*x, a*) through the identity:

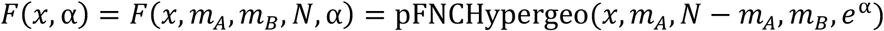

The test-based confidence interval *CI*(*X*, 1 – *γ*) defined above in *(4)* is precisely the interval CI.CP, labeled CP because of the close resemblance of this test-based idea to that of Clopper and Pearson (Clopper & Pearson, 1934) in a famous 1934 paper about confidence intervals for binomial proportions. By definition, it is *conservative* in the sense that its probability of covering the true *α* value generating the random variable *X* is at least 1 – *γ*. However, due to the discreteness of the Extended Hypergeometric distribution, for moderate *m_A_*, *m_B_*, *N* the excess of the actual coverage probability over the nominal 1 – *γ* can sometimes be quite large, up to 0.03 or more when *γ* = 0.05.

#### Blaker Confidence Interval

Another type of confidence interval defined by (Blaker, 2000) is proved at the same time to be conservative and also a subset of CI.CP, and is found for some combinations of (*α*, *X*) to improve the coverage probability substantially. Defining and justifying the construction of Blaker Confidence Interval (CI.Blaker) is a little more abstract and difficult, and we refer the interested reader to the Splus code and Theorem 1 of (Blaker, 2000), defined in that paper for unknown Binomial proportions, that we have adapted in our package functions `AcceptAffin` and `AcceptAffCI`.

The coverage performance of the CI.CP in the context of Binomial proportions, was compared by (Brown, Cai, & DasGupta, 2001) with several other popular confidence intervals, mostly inspired by the large-sample closeness (the DeMoivre-Laplace Central Limit Theorem) of the binomial to the normal distribution with the same mean and variance. The CI.CP was found to be extremely conservative, and several well-known and widely used large-sample-theory-based intervals were found to have (for some (*n, p*) combinations) extremely erratic performance, for surprisingly large values of *n*. (That observation, and some theoretical explanation of it based on deeper large-sample theory, was the main point of the (Brown et al., 2001) article and other theoretical articles on which it was based.) Although that study compared several confidence intervals, it did not study the modified Clopper-Pearson intervals known as midP (CI.midP) [(Agresti, 2013), p. 605]. Both the Clopper-Pearson and midP intervals are applicable to small- and as well as large-sample data and have natural generalizations to unknown scalar parameters for other discrete random variables like the Extended Hypergeometric.

#### Preference for Conservative or Close-to-nominal-coverage Intervals?

Should confidence intervals always be conservative to be useful? For some applications, such as the confirmatory clinical trials of governmental statistical regulatory agencies (FDA, NIH, NIST) and some laboratory testing, the answer is yes. But for a great deal of exploratory work in science, including ecology, it will often be preferable to choose a type of confidence interval with coverage probability reliably as close as possible to the nominal coverage 1 – *γ*, even at the cost of occasional under-coverage by a few percent. With this in mind, we present two other confidence intervals for the log-affinity parameter *α*, the midQ and midP intervals, which use the same strategy as CI.CP but calculate interval endpoints not from distribution-function values that jump at each successive integer co-occurrence value, but respectively with linearly interpolated inverse distribution function values between successive integers or linearly interpolated distribution function values. As it turns out, these two approaches give nearly identical results (see below).

#### midQ Confidence Interval

First, as we explained in “Advances: The median interval”, we define the interval of all *α* values for which the observed co-occurrence count to *X* is the *q*’th quantile for *F*(*x, m_A_, m_B_, N*, α) as

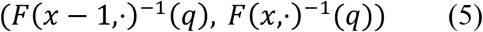

To estimate a single *α* value for which *X* is a *q*’th quantile, we choose the midpoint of the interval *(5)*. Then a natural choice for an interval of *α*’s compatible with and observed cooccurrence count *X*, expressed in terms of the quantiles *γ*/2 and 1 – *γ*/2, is the interval

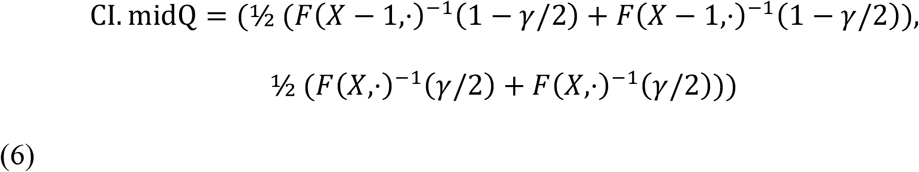

This interval has coverage probability closer to the nominal level 1 – *γ* than the conservative intervals CI.CP and CI.Blaker described above, at the cost of occasional undercoverage (e.g., by as much as 0.03 for 95% intervals, for some specific combinations of *α* and *m_A_, m_B_*, *N*).

#### midP Confidence Interval

Another confidence interval Midpoint Exact or MidP (CI.midP), which has almost exactly the same behavior for most (*X, m_A_, m_B_*, *N*) as CI.midQ, is defined directly in terms of approximate distribution function values rather than quantiles. This idea, which gets the midP name from the analogous interval for unknown binomial proportions [(Agresti, 2013), p. 605], is the interval defined exactly like CI.CP with the distribution function *F*(*x*, *α*) replaced by ½(*F*(*x* – 1, *α*) + *F*(*x*, *α*)):

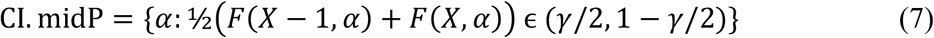

The intervals given so far are defined only for *X* values away from the extremes of its range. When *X* = max(*m_A_* + *m_B_ – N*, 0), the left endpoint of the confidence interval should be −∞, except that we replace that value by −log(2*xN*^2^), which was mentioned earlier to be a provable lower-bound for all 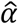 corresponding to *X* > max(*m_A_* + *m_B_* – *N*, 0). Similarly, when *X* = max(*m_A_, m_B_*) the right endpoint of all CI’s is taken to be log(2*N*^2^). One further modification in the `CooccurrenceAffinity` package, adopted to preserve precision of the CI’s computed from the `BiasedUrn` package evaluations of *F*(*x*, *α*), is the restriction that all confidence intervals are replaced by their intersection with the interval (–*M, M*), where *M* denotes the value at which we earlier mentioned that 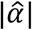 is capped.

#### Good practice

The theoretical discussion of confidence intervals for *α* presented under “Advances: Four types of confidence interval” and their examples below may be useful in setting some direction/recommendation about which CI to use. In confirmatory applications of confidence intervals for which strict control of the type I error probability is desirable, we recommend CI.Blaker. By contrast, in exploratory studies we may be comparing many confidence intervals for different datasets and might for that reason want type I error probabilities (which equate to probabilities of not covering the true *α*) to be as close as possible to the usual nominal non-coverage, usually 0.05. In such studies, we recommend either of the nearly indistinguishable midP or midQ intervals.

## OVERVIEW OF CooccurrenceAffinity PACKAGE

The functions of this package are visually shown in Fig. 3 based on their role (color coded) and in relation to each other (indicated by the proximity and arrow). Several of these functions rely on the output of another R package `BiasedUrn`.

**Figure 3.**
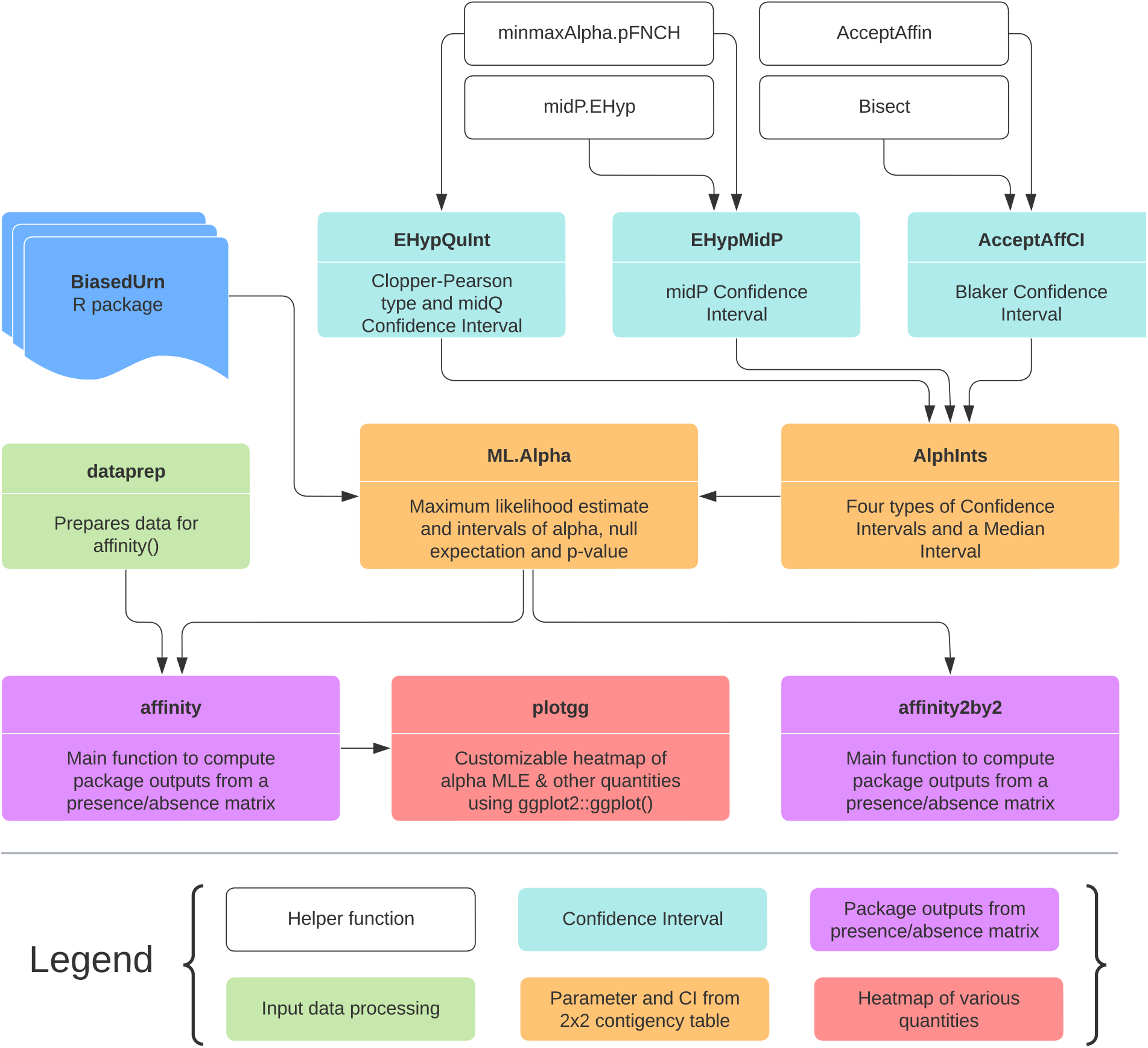
Workflow of `CooccurrenceAffinity`. The functions are grouped under different colors based on their role indicated in the box. A function at the base of an arrow feeds its output to the function at the tip of the arrow. Not all functions of the package are shown here.

### Functions to compute four types of confidence intervals

`EHypQuInt` computes Clopper-Pearson type Confidence Interval and midQ Confidence Interval. `AcceptAffCI` and `EHypMidP` compute Blaker Confidence Interval and midP Confidence Interval, respectively. These three functions rely on the following four helper functions for computation of confidence intervals: `minmaxAlpha.pFNCH, AcceptAffin, Bisect`, and `midP.EHyp`.

### Function for parameter estimate and all intervals (median and CI)

`AlphInts` lists all the four confidence intervals and computes median interval. It also reports null-distribution expected co-occurrence count and p-value. This function feeds all of its outputs to `ML.Alpha` which additionally computes the parameter estimates 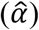, maximized log-likelihood.

### Functions for analysis of 2×2 contingency table

`AlphInts` and `ML.Alpha` can be used for cooccurrence analysis based on a 2×2 contingency table show in Fig. 1. However, if the goal is also to obtain the values for the common traditional indices, `affinity2by2` can be useful. `affinity2by2` supplements the outputs of `ML.Alpha` with the indices of Jaccard, Sørensen–Dice, and Simpson. These indices are useful for reference to older studies but should be used with caution.

### Function for analysis of presence-absence matrix

A presence-absence matrix (e.g., species by site matrix) is analyzed with `affinity`. The input dataset is examined for any potential issues and a ready-to-analyze matrix is prepared by calling a separate function `dataprep`. A user should indicate if rows or columns are being analyzed. In a species by site matrix, if species are given in rows and islands in columns, an analysis of rows gives affinity between every pair of species whereas an analysis of columns gives affinity between every pair of islands. A user can also optionally select certain rows or columns based on their position or name. This function accepts abundance data, in which case, a user should indicate datatype = “abundance”, and should pick a threshold to convert abundance data to binary one. The user should also indicate if the threshold should be considered in the category of presence or absence with “class0.rule” argument.

`affinity`, by default, returns two matrices. The first one is a long-format data frame with each pairwise analysis on a row and a total of 19 columns for various outputs of analysis. The second matrix is the processed presence-absence matrix used for the analysis (e.g., binary version of the abundance matrix, a subset of the original matrix with selected rows/cols). `affinity` optionally returns (with “squarematrix” argument) up to 11 additional square matrices for 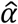 and other quantities. A square matrix holds the entities being analyzed both on the row and columns in the same order and each cell of the matrix has the values of a particular quantity (e.g., 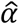).

### Function to plot the output

`plotgg` generates a heatmap for any quantitative column in the long-format output available under `$all` of `affinity` output. `plotgg` utilizes `ggplot2::ggplot` on the backend. The arguments of `plotgg` make it easy to customize the plot.

### Function to test true coverage of confidence intervals

`CovrgPlot` computes the true coverage probabilities of the selected confidence intervals and returns, for each confidence interval type, a multi-panel plot with (a) line plot of the coverage probabilities by 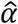, and (b) a histogram of the coverage probabilities, showing percentage of the probabilities falling below (rate of failure to include population parameters) *vs*. above the confidence level. Following a plan similar to that described computationally in (Brown et al., 2001) regarding confidence intervals for Binomial proportions, our package function `CovrgPlot` calculates and plots the curve of coverage probabilities under the Extended Hypergeometric distribution for fixed (*X, m_A_, m_B_, N*) and *all α* values.

`Covrg` is a way to generate the set of 4 coverage probability values without plotting anything. The coverage probabilities were calculated (and kept as an output array) in `CovrgPlot` at values where the coverage curve changes direction, but can be separately calculated at any single *α* using `Covrg`.

## EXAMPLES AND ILLUSTRATIONS

### Analysis of 2×2 contingency table

We compute with a running example *X*=35, *m_A_*=50, *m_B_*=70, *N*=150. The syntax and results of the function calls for figuring the MLE 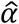, the median interval, and the 95% two-sided equal-tailed confidence intervals for *α*, are shown in Fig. 4.

**Figure 4.**
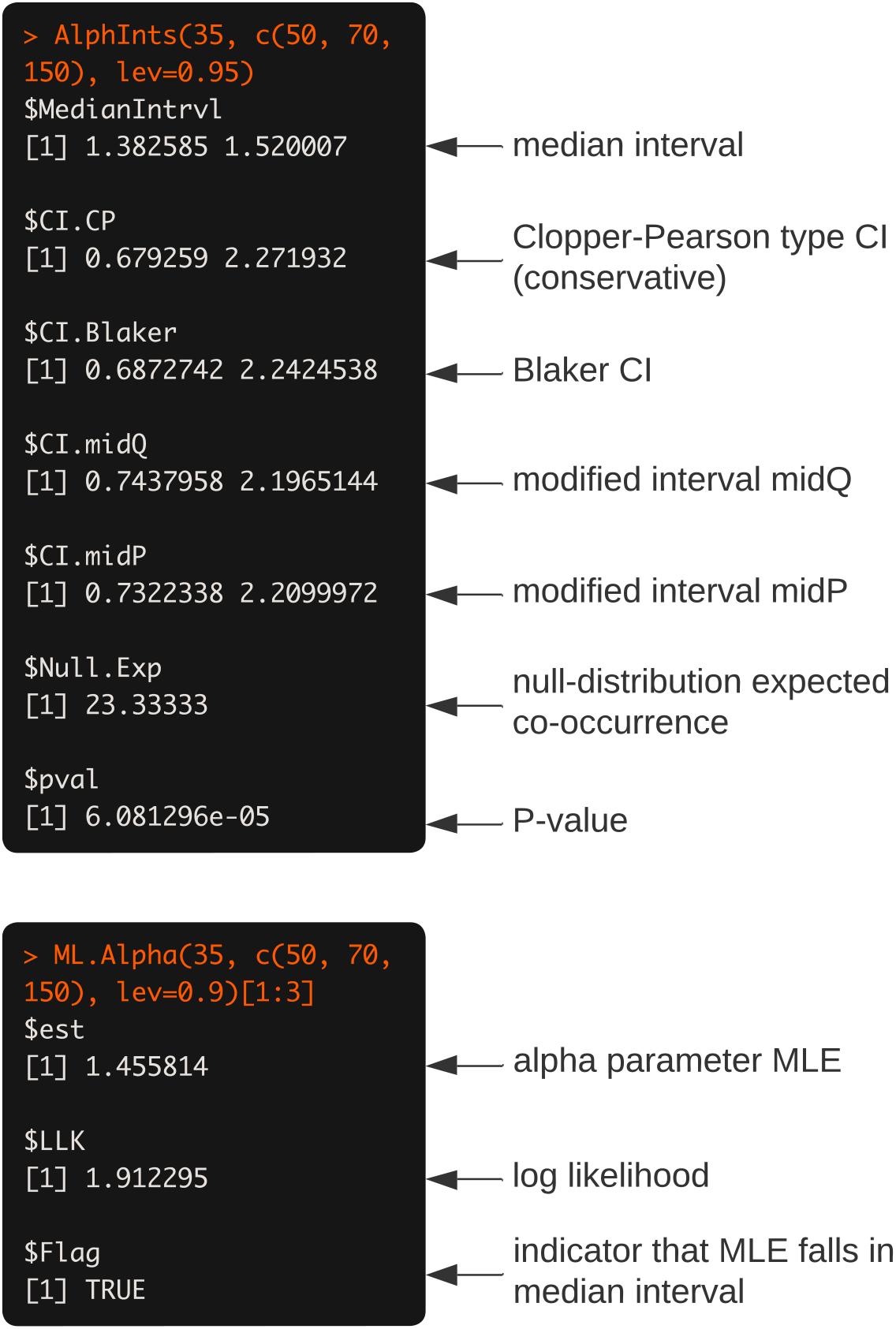
An example of syntax and output of `AlphInts` and `ML.Alpha`.

The conservative interval of Clopper-Pearson type CI is definitely wider than other intervals (Blaker, midQ and midP). The MLE computed here is very close to the log cross-product ratio, which is also what the `loglin` base-package loglinear-modeling function gives, as in Fig. 5.

**Figure 5.**
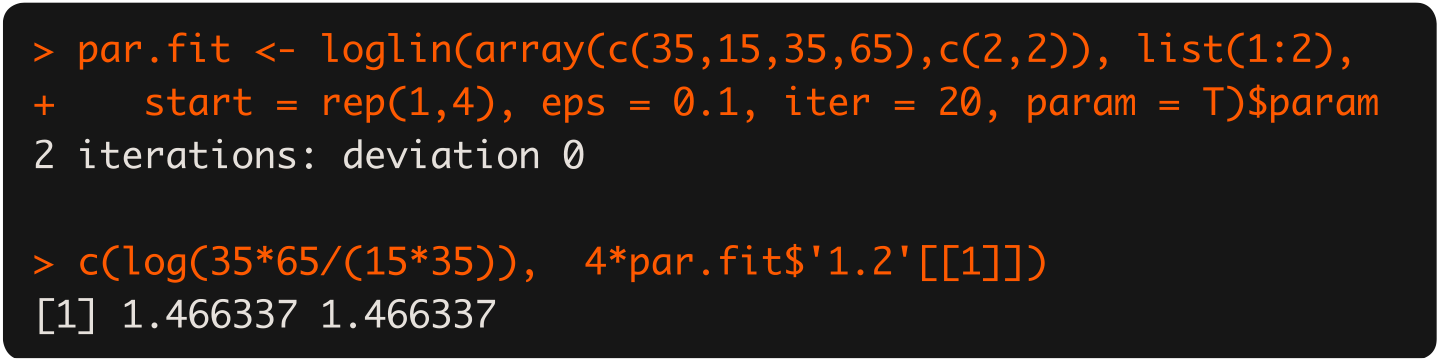
The log cross-product ratio of the example in Fig. 4.

As we saw above, the MLE 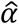 in this instance is 1.455814. The difference between this estimate and the `loglin` output is that here we condition on the marginals while `loglin` does not. Another method of computing MLEs is Poisson regression, which also does not condition on marginals and gets the same answer as `loglin`. Especially in small to moderate samples, (Mainali et al., 2022) argue that it is important for the index 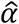 to be insensitive to prevalences *m_A_, m_B_*. Therefore, the MLE conditioned on marginals makes the most sense for ecological analyses.

### Analysis of presence-absence matrix

For users with presence-absence matrix interested in several pairwise analysis, affinity is more useful. For each pair of entities, affinity computes 2×2 contingency table from presence-absence data and performs the analysis. An example showing computation of affinity between species or between sites from the same dataset is presented in Fig. 6.

**Figure 6.**
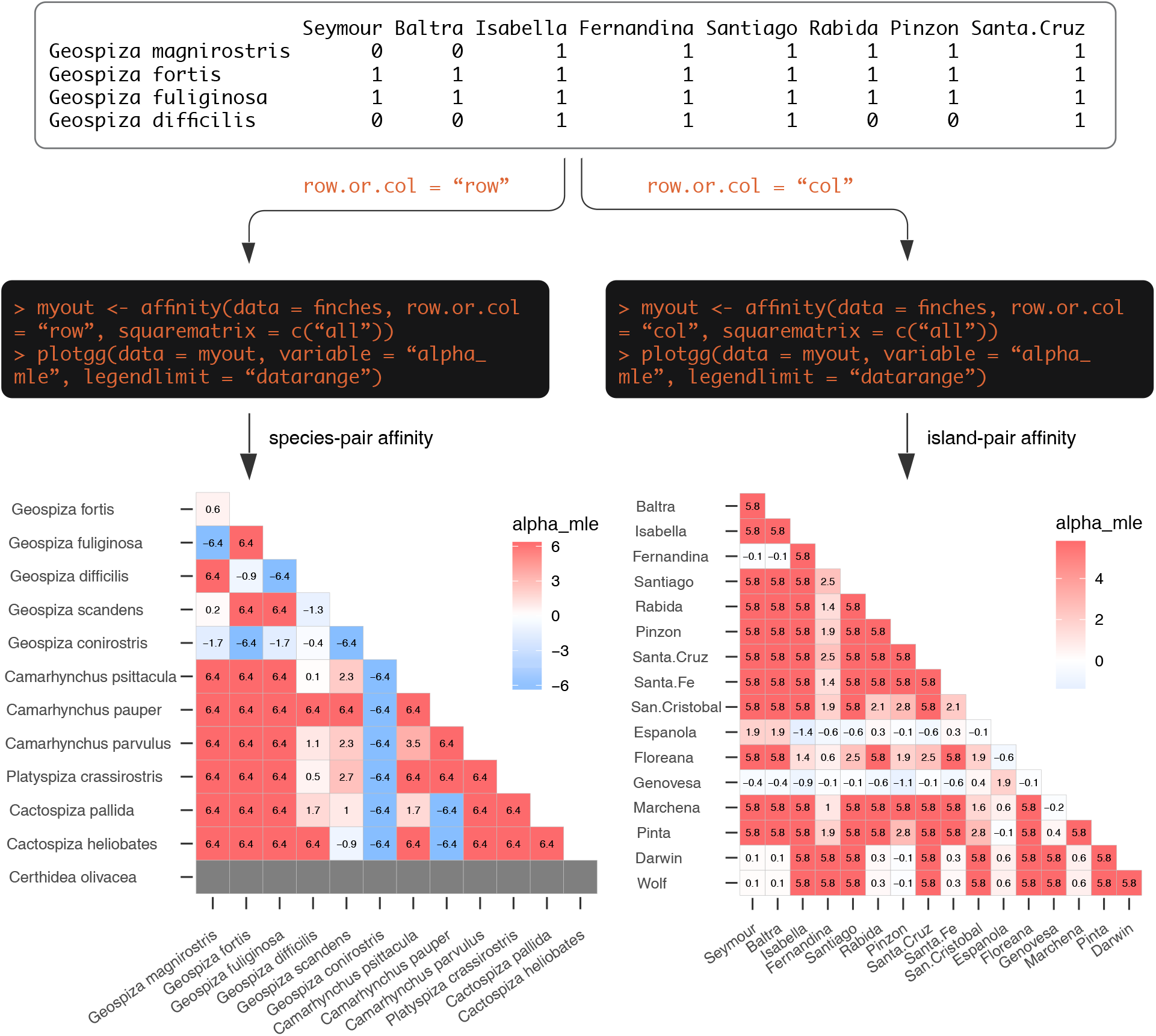
Analysis of a presence-absence matrix for computing several pairwise analysis for row or column. The top panel shows part of the data. Code block shows how easily affinity between species (rows) or between islands (columns) can be computed with `affinity` and the results can be plotted with `plotgg`.

### Quantifying discreteness versus coverage

The median interval width is a measure of the discreteness of the cooccurrence distribution, while the confidence intervals have the usual interpretation of confidence intervals. The practical interpretation is that the CI’s are imprecise (due to discreteness) roughly by the magnitude of median-interval width.

We illustrate an example of the relative sizes of the median interval and confidence interval and their positioning with respect to the MLE (Fig. 7). Examining the trend for all possible values of *X* from 1 to 49, we can see that the interval estimates of median interval and Blaker CI tend to become wider for more extreme co-occurrence values of *X*, while the median interval is always much shorter than the Blaker CI (Fig. 7a, other CIs not shown). Blaker CI and Clopper-Pearson type CI tend to be very similar in length and positioning when the cooccurrence count is near its null expectation, while the Blaker CI tends to be shorter relative to Clopper-Pearson type CI for *X* values farther from that null expectation, which is 50*70/150 = 23.3 in the Fig. 7b.

**Figure 7.**
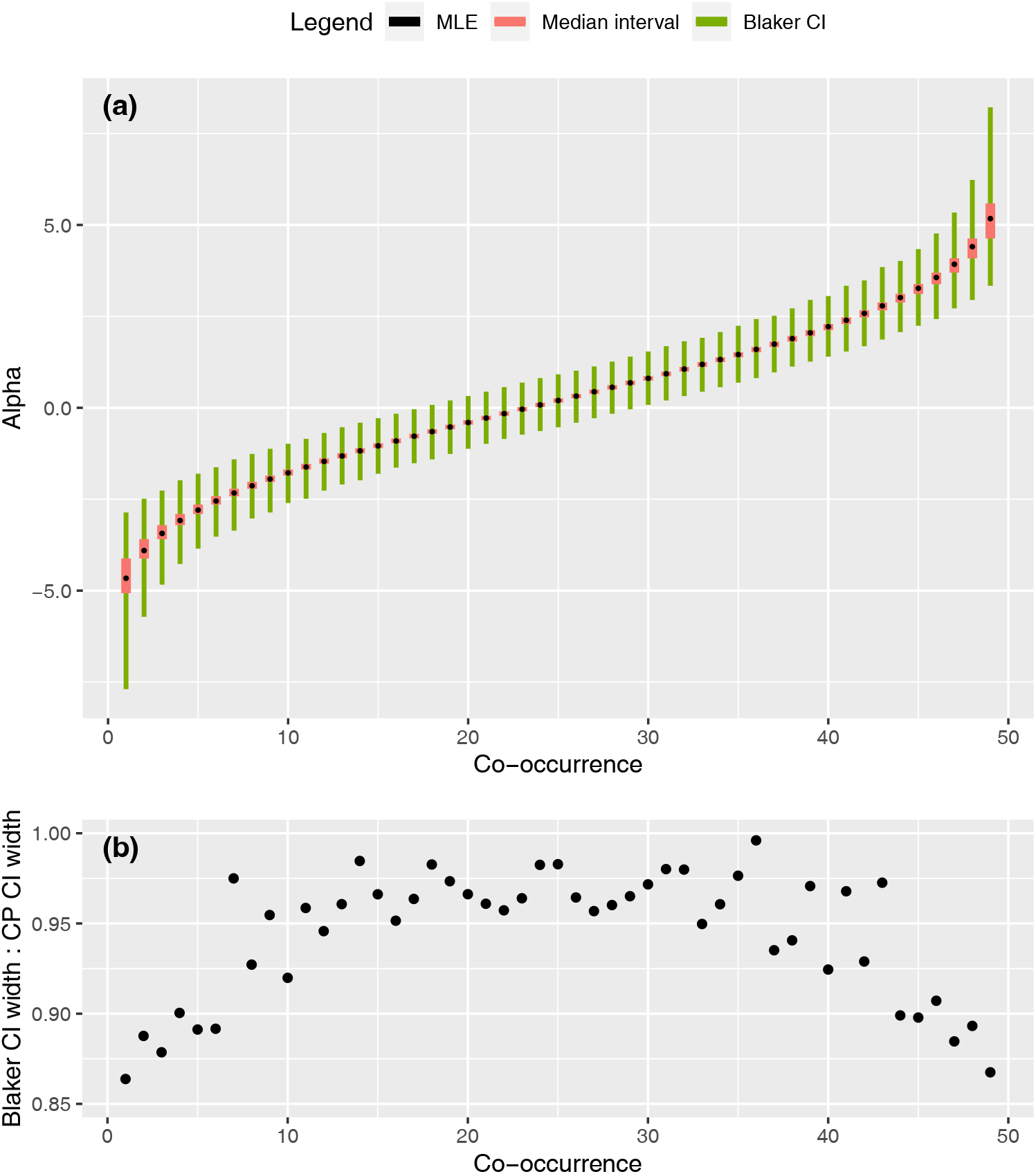
Intervals and MLE for *α* for all possible *X* values (co-occurrence) in an example of *m_A_* = 50, *m_B_* = 70, *N* = 150. (a) MLE, median interval and Blaker CI (95%), (b) ratio of the length of Blaker CI (95%) to that of Clopper-Pearson type CI (95%).

A further picture to clarify the construction of the confidence intervals is given in Fig. 8, showing the function values *F*(*x*, *α*) for various choices of *x*. Note that the values *F*(*x, α*) decrease as a function of *α*, because larger *α* corresponds to larger co-occurrence random variable *X* and smaller probability for that *X* to be ≤ *x*. As an example of the information to be read from the figure, for X=20: the median-interval in green horizontal bar is the alpha-interval between the pair of green curves at height *F*(*X*) = 0.5, with 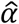 the alpha-coordinate of the solid black circle between the two curves. The Clopper-Pearson confidence interval has left endpoint at *α* where the left green curve crosses level 0.975 and right endpoint where the right green curve crosses level 0.025.

**Figure 8.**
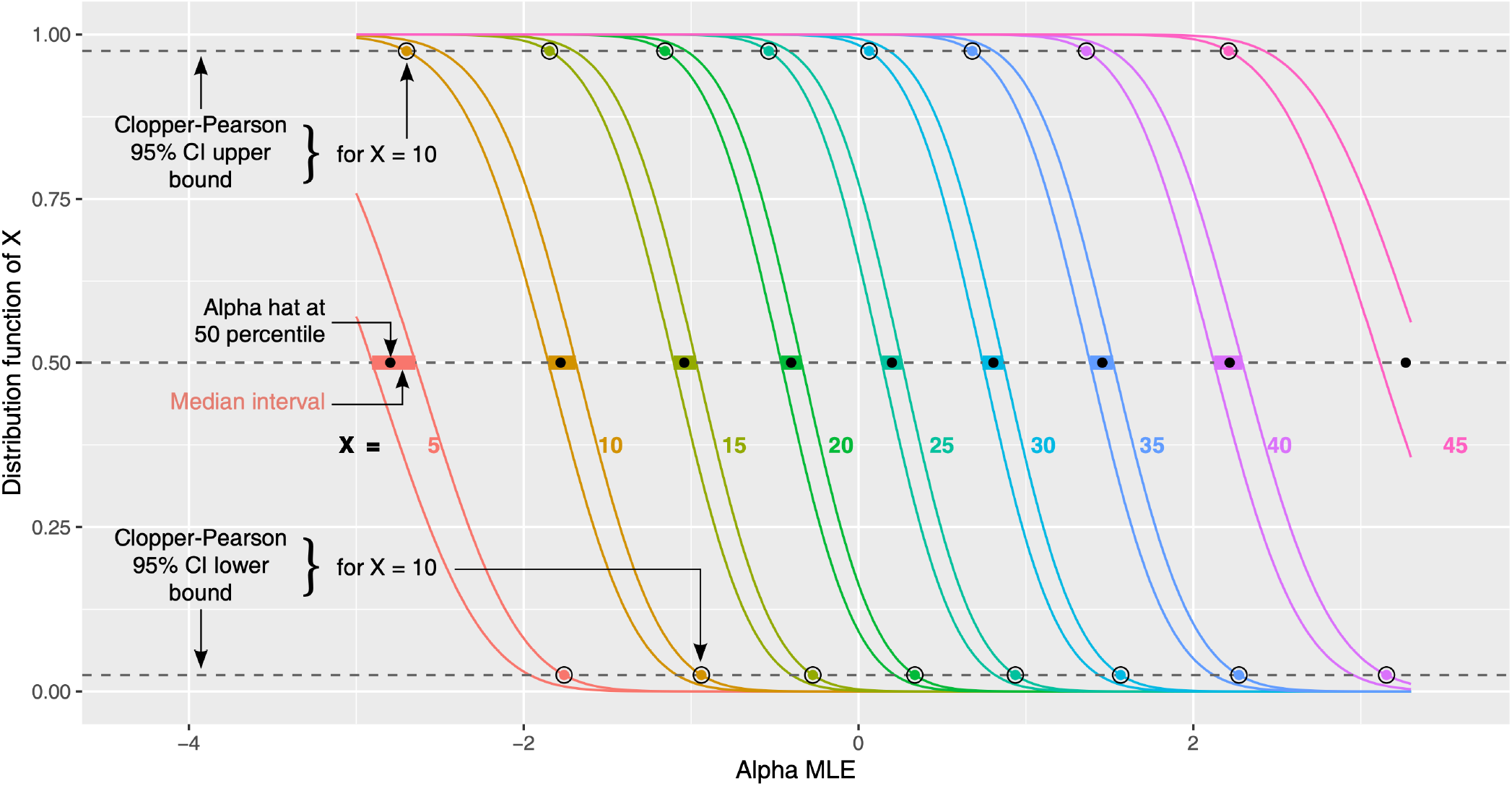
Functions *F*(*x, α*) plotted for selected pairs *x, x* – 1 and all *α* in the range (−3, 3.3) for the same example of *m_A_* = 50, *m_B_* = 70, *N* = 150. The colored sigmoid lines are the curves *F*(*X*,·), with *x* indicated in the same color, and immediately to the left of each such curve is another (with the same color) for *x* – 1. Horizontal dashed lines are the quantile levels 0.025, 0.50, 0.975. Black solid circles are plotted for the *X* values in each curve-pair at (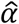, 0.5). Colored solid circles encircled by black circles are plotted for the upper and lower bounds of Clopper-Pearson 95% confidence intervals. The median-interval of *α* ‘s for each *X* is the colored horizontal bar connecting the curve-pair at height 0.5.

The four types of CIs differ in their span, which has implications for their true coverage probability. The true coverage probability is the probability that the interval endpoints bracket the true population parameter. Ideally, the probability should be equal to the confidence level, but this is not quite true for discrete data. By inspecting the trend of probabilities, we can see several patterns (Fig. 9). First, the true coverage probability of including population parameter by 95% Clopper-Pearson type CI (Figs 9a,b) and Blaker CI (Figs 9c,d) is always 95% or more (histograms show 100% of the mass above the 0.95 probability). In order to minimize the rate of failing to include the population parameter, these two CIs become too wide. However, the Clopper-Pearson CI interval is (unnecessarily) wider than the Blaker CI interval; the latter is closer (within 0.025) to the correct coverage compared to the former (within 0.05). Second, the 95% interval of midP (Figs 9e,f) and midQ (Figs 9g,h) are narrower than the other two; the true coverage probabilities of midQ and midP tend to center around 0.95 for the range of *α* values seen with all possible cooccurrences. Roughly half the time (53.59% for midQ and 45.3% for midP, shown in histograms), these CIs have actual coverage probability below (but within 0.02 of) the nominal value 0.95.

**Figure 9.**
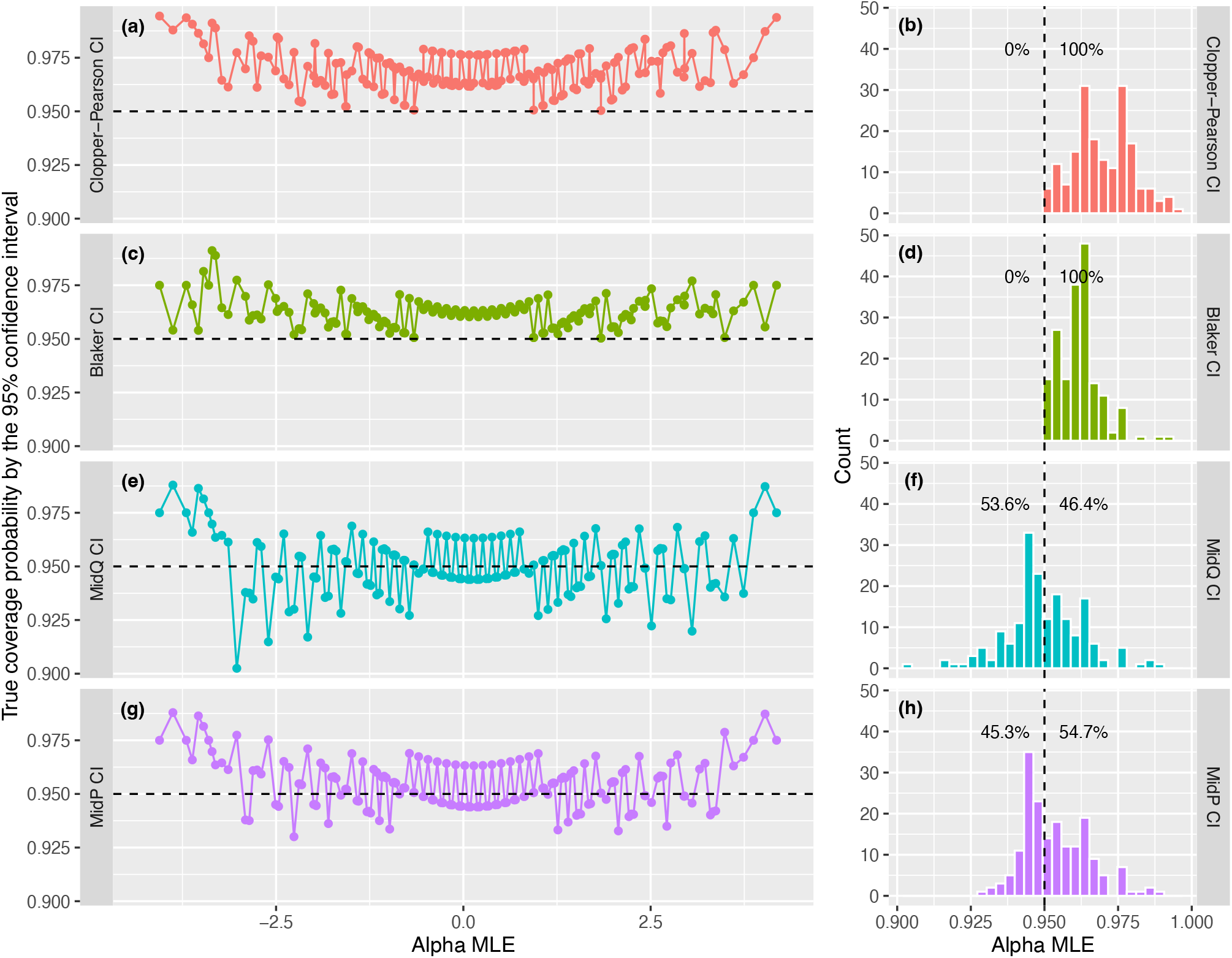
True coverage probability of the four types of confidence intervals for the example presented in Fig. 7 with *α* values shown from −4 to 4, computed as `CovrgPlot(c(50,70,150), lev=0.95)`. The probability of including population parameter (true coverage probability) by 95% CI of Clopper-Pearson type CI is always at least 95% (dashed line) for all co-occurrences (a) resulting in a histogram with its entire mass above 0.95 (b). Such pairs of plots are also shown for Blaker CI (c,d), midQ (e,f) and midP (g,h) CIs.

The coverage of the midP type interval is close (within ±0.02) to the correct coverage and averages across *α* to almost exactly the nominal 95%, while the Clopper-Pearson type interval is conservative by as much as 0.05. This picture relates to only one (*m_A_, m_B_*, *N*) combination, but is the result of a rapid exact calculation, not a simulation, and so can easily be reproduced for other fixed margin values.

## CONCLUSION

`CooccurrenceAffinity` is a new R package that computes parameters associated with a recently developed metric of association in co-occurrence data (Mainali et al., 2022). This novel metric, called alpha, corrects for the pervasive errors in analysis of co-occurrence data with traditional indices. This package includes flexible functions for (1) analyzing 2×2 contingency tables of count of occurrences or a more widely used m×n matrix of presence-absence, and (2) plotting the analysis output. The novel elements of this paper are the introduction of median interval and four types of confidence intervals of alpha. The package provides functions to (a) compute these intervals, and (b) evaluate their efficacy of true coverage probability of including population parameter, and (c) visually and numerically evaluate the balance between their rate of failing to include the population parameter α and their width.

We expect that this package will serve as a user-friendly tool for ecologists and biogeographers to estimate affinity in co-occurrence data. Given the current surge in environmental and spatial dataset integration in cloud computing environment where Python is emerging as the first choice for data science, a Python adaptation of the package would be helpful to the communities.

## ACKNOWLEDGEMENTS

KM was supported by the Grayce B. Kerr Fund, Inc, and by the National Science Foundation DBI-1639145 under funding received for the National Socio-Environmental Synthesis Center (SESYNC).

## AUTHORS’ CONTRIBUTIONS

ES and KM wrote the R functions; KM developed the GitHub package; KM and ES wrote the vignette; KM and ES conducted the package tests; ES and KM wrote the package website and function documentations; KM and ES wrote the manuscript.

## DATA AND PACKAGE AVAILABILITY STATEMENT

The package introduced in this publication is available at the following GitHub repository: <https://github.com/kpmainali/CooccurrenceAffinity>. This repository carries R functions and data used in this publication, package documentations and vignettes. The actual R script used to generate figures is included in the figures themselves or in a separate GitHub repo < https://github.com/kpmainali/CooccurrenceAffinity_PublicationScript>.

